# Climatic predictors of prominent honey bee (*Apis mellifera*) disease agents: *Varroa destructor*, *Melissococcus plutonius*, and *Vairimorpha* spp

**DOI:** 10.1101/2024.06.28.601058

**Authors:** Alison McAfee, Niloofar Alavi-Shoushtari, Lan Tran, Renata Labuschagne, Morgan Cunningham, Nadejda Tsvetkov, Julia Common, Heather Higo, Stephen F. Pernal, Pierre Giovenazzo, Shelley E. Hoover, Ernesto Guzman-Novoa, Robert W. Currie, Patricia Wolf Veiga, Sarah K. French, Ida M. Conflitti, Mateus Pepinelli, Daniel Borges, Elizabeth M. Walsh, Christine A. Bishop, Amro Zayed, Jason Duffe, Leonard J. Foster, M. Marta Guarna

## Abstract

Improving our understanding of how climate influences honey bee parasites and pathogens is critical as weather patterns continue to shift under climate change. While the prevalence of diseases vary according to regional and seasonal patterns, the influence of specific climatic predictors has rarely been formally assessed. To address this gap, we analyzed how occurrence and intensity of three prominent honey bee disease agents (*Varroa destructor* ― hereon *Varroa* ― *Melissococcus plutonius*, and *Vairimorpha* spp.) varied according to regional, temporal, and climatic factors in honey bee colonies across five Canadian provinces. We found strong regional effects for all disease agents, with consistently high *Varroa* intensity and infestation probabilities and high *M. plutonius* infection probabilities in British Columbia, and year-dependent regional patterns of *Vairimorpha* spp. spore counts. Increasing wind speed and precipitation were linked to lower *Varroa* infestation probabilities, whereas warmer temperatures were linked to higher infestation probabilities. Analysis of an independent dataset shows that these trends for *Varroa* are consistent within a similar date range, but temperature is the strongest climatic predictor of season-long patterns. *Vairimorpha* spp. intensity decreased over the course of the summer, with the lowest spore counts found at later dates when temperatures were warm. *Vairimorpha* spp. intensity increased with wind speed and precipitation, consistent with inclement weather limiting defecation flights. Probability of *M. plutonius* infection generally increased across the spring and summer, and was also positively associated with inclement weather. These data contribute to building a larger dataset of honey bee disease agent occurrence that is needed in order to predict how epidemiology may change in our future climate.

## Introduction

Infectious diseases and parasites are among the dominant factors affecting honey bee (*Apis mellifera*) colony health in commercial operations.^1–3^ Several recent studies have investigated relationships between climatic variables and overwintering colony mortality,^4–8^ but similarly large-scale association studies between weather patterns and specific diseases or pathogens are more rarely conducted, with the notable exception of Rowland *et al.,*^9^ who examined patterns of disease incidence associated with climatic variables in England and Wales. Weather patterns are expected to become less stable as the climate changes, with higher frequencies of extreme events along with a shifting baseline to warmer temperatures and region-specific shifts in precipitation.^10–12^ A better understanding of relationships between honey bee disease agents and current climatic trends would be an asset for predicting how these agents might change in the vicissitudes of our future climate.

Several etiological agents are known to occur in seasonal cycles, including but not limited to *Vairimorpha* spp. (formerly *Nosema* spp.,^13^ the causative agent(s) of nosemosis), *Melissococcus plutonius* (the causative agent of European foulbrood disease, or EFB), and the mite *Varroa destructor* (hereon referred to as *Varroa*, the causative agent of varroosis).^14–21^ Such seasonal patterns may be driven by direct effects of weather on pathogen or parasite survival and reproduction, or indirect effects via associated changes within the honey bee colony. As an example, high temperatures (>35°C) reduce *Varroa* survival and reproduction directly,^22,23^ whereas moderately increasing temperatures during the spring lead to increased honey bee brood rearing, which *Varroa* in turn requires for reproduction.^14^ Moreover, favorable conditions (warm temperatures with low wind and precipitation) increase dispersal opportunities between colonies for disease agents, but might also affect the risk of colonies manifesting diseases that are associated with poor food availability or stress (such as EFB^24^). It is thus difficult to predict, *a priori*, how weather conditions may translate into changes in intensity or occurrence of disease agents.

*Varroa* is broadly considered the most economically damaging parasite of honey bees, and its life cycle is closely tied to seasonal changes in honey bee brood production.^14,25^ The mites disperse to new colonies through contact between bees at forage sources as well as robbing and drift (see reviews^14,25^ for more information about *Varroa* biology and control). *Varroa* has successfully become established in all continents where honey bees are maintained, and only a handful of island territories remain *Varroa*-free.^14,26^ Many studies broadly highlight the impact of winter brood breaks on *Varroa* loads and differences in *Varroa* population growth in temperate versus tropical climates (reviewed in Rosenkranz et al.^25^). The studies that have investigated relationships with weather conditions have focused mainly on temperature,^9,22,23,27^ with extreme heat being associated with reduced *Varroa* reproduction^22,23^ but otherwise a positive relationship between *Varroa* occurrence and temperature has been observed.^9,27^ One study conducted by Harris *et al.,*^23^ found that rainfall had no association with *Varroa* population growth rates, and a recent study by Rowland *et al.*^9^ found a negative correlation between *Varroa* occurrence and precipitation and wind. Further studies in different regions would be helpful to clarify and substantiate these findings in different geographies.

*Vairimorpha* spp. are microsporidian endoparasites of the honey bee midgut and, in temperate climates, tend to infect bees with seasonal intensities that are normally highest in the spring and early summer, then decrease in late summer and early fall.^18–21^ Two species of *Vairimorpha* are known to infect honey bees in Canada: *V. apis* and *V. ceranae*. with different seasonal trends. Whereas *V. apis* exhibits a regular peak in average gut spore loads in the spring and often a secondary peak in in the fall,^28^ *V. ceranae* trends appear more variable across different countries.^29–32^ Regional differences in *V. ceranae* and *V. apis* abundance has led to speculation over whether each species’ relative success is influenced by climate, but it is not clear if this is the case.^17,19^ *V. ceranae* is the dominant species in Canada and the United States;^33,34^ therefore, in these regions detection metrics that do not discriminate between species (*e.g.* spore counts) likely mainly represent *V. ceranae.* Some^18,35^ but not all^36^ field studies have found that higher temperatures correlate with lower spore loads, and some research suggests that geographic region (and, by extension, climate) is an important factor influencing *Vairimorpha* spp. epidemiology.^37^ Indeed, in addition to warm temperatures inhibiting *V. apis* and *V. ceranae* proliferation in laboratory trials,^38^ inclement (cold, windy, or wet) weather limits opportunities for defecation flights that help clear *Vairimorpha* spp. infections,^39,40^ but interactions among these variables have, to our knowledge, not yet been investigated.

EFB disease can cause severe damage to honey bee colonies by killing developing brood, thereby weakening the colony and reducing productivity.^16^ *M. plutonius* is often detected in asymptomatic and apparently healthy colonies, especially those proximal to symptomatic colonies and apiaries;^16,24,41^ therefore, dispersal from infected units is likely an important factor governing patterns of *M. plutonius* prevalence. EFB is thought to be an opportunistic disease that tends to manifest when larvae are under nutritional stress, such as when colonies are producing new brood very quickly in the spring or when pollen and nectar resources are low.^16,24^ However, larval food supply is clearly not the only factor involved, since larvae fed in excess may still develop the disease.^42^ Interestingly, Milbrath^16^ and Bailey^43^ suggest that even favorable foraging conditions could lead to temporary nutritional stress of larvae during the spring if a strong nectar flow causes nurse bees to reallocate their labor to process honey. Rowland *et al.*^9^ found that higher precipitation (associated with poor foraging conditions) was linked to higher EFB incidence in England and Wales, but no relationship was found with temperature nor wind speed.

Given how widespread these honey bee pathogens and parasites are and their apparent associations with weather conditions, we sought to investigate relationships between the disease agents and climatic variables while accounting for broad differences between regions and years. To this end, we used a dataset of *Varroa* infestation levels, *Vairimorpha* spp. spore intensities, and *M. plutonius* occurrence across five Canadian provinces, representing a wide range of climates and geographies, to identify associations with temperature, precipitation, and wind speed. While regional differences were most pronounced, climatic variables were also influential and largely reveal intuitive patterns. This dataset will help build the growing body of knowledge and data necessary to facilitate predictive modelling of disease agent dynamics under projected climate conditions.

## Methods

### Honey bee colonies and datasets

This study includes analyses of a primary dataset (*Varroa*, *Vairimorpha*, and *M. plutonius* data collected in 2020 and 2021) and a validation dataset (*Varroa* only, collected in 2016). Some, but not all, elements of the primary and validation datasets have been previously described in several publications (see details below).

A subset of the primary dataset’s honey bee colonies and pathogen/parasite data have been previously described by French *et al.*^44^ (240 colonies sampled in 2020 and 2021) along with additional colonies (a further 240 in both years) described here and partially reported in McAfee *et al*.^45^ As described further below, samples for *M. plutonius* and *Vairimorpha* spp. testing were pooled ahead of analysis (4 colonies per composite sample), whereas samples taken for *Varroa* detection were not pooled. The total number of samples analyzed for *M. plutonius*, *Vairimorpha* spp., and *Varroa* were therefore N = 120, 120, and 480, respectively, though all were derived from the same 480 colonies.

As previously described,^44^ the colonies were located in five provinces (British Columbia (BC), Alberta (AB), Manitoba (MB), Ontario (ON), and Quebec (QC)), and eight different regions. Within these regions, colonies were derived from the same beekeeping operation; therefore, operational and regional differences are indistinguishable. Two regions/operations were represented within AB (near Grande Prairie to the north and Lethbridge to the south); three regions/operations were represented in QC (near Quebec City in 2020 and near Montreal and Lac St. Jean in 2021); and BC, MB, and ON were represented by one region/operation each. Colonies were placed near or far from one of eight possible focal crops, which depended on region, and sampled at the start (immediately before or immediately after moving into pollination), middle (peak bloom), and end (immediately before or immediately after) of the pollination period, the duration of which varied depending on commercial standards for each crop. In the case of corn, which is not pollinated by honey bees, colonies were sampled before sowing, during sowing, and at the end of the growing season. An equal number (N = 240) of additional colonies produced from the same stocks, according to the same standards, as those assigned to the focal crops but located far from the pollination yards (>1.5 km, with the exception of highbush blueberries which were > 1.3 km) are included in this dataset, bringing the final number of colonies to 480 across site types (near or far from crops) in both years, each subject to three sampling events over time. A subset of these data are also described in McAfee *et al*.^45^ Colony locations in each province and at each time point are illustrated in **Supplementary Figure S1**.

In each region, colonies were either supplied by collaborating beekeepers, owned by the relevant institution, or purchased from producers, but in all cases sampling and management during the experiment were conducted by the research teams. As previously described,^44^ colonies were headed by locally overwintered queens and colony sizes were standardized in deep Langstroth-style single-or double-box pollination units according to standards for commercial pollination for each focal crop. Although *Varroa* is one of the parasites of interest, which we subsequently measured, early season *Varroa* control is often necessary to ensure colony vitality throughout the season; therefore, in some cases colonies were treated with miticides before the experiment began if adult bee infestation levels exceeded the economic threshold (>1 mite per 100 bees).^46^ Specifically, in early spring, colonies in BC (2020; n = 64) were treated with Formic Pro and colonies in southern Alberta (2020 and 2021; n = 48 in both years) were treated with Apivar according to manufacturer instructions. Mite treatments applied to colonies prior to being purchased from commercial suppliers (BC in 2021 [n = 80] and MB in 2020 and 2021 [n = 56 and 16, respectively]) are unknown. All other colonies in northern AB (n = 16 in both years), ON (n = 40 in 2020), and QC (n = 16 in 2020 and 80 in 2021) were under university or government research laboratory management for the full calendar year and miticides were not applied. These differences in treatment (and other operational/regional differences) were accounted for statistically by including region as an interactive term in the model and by including colony as a random intercept term (accounting for differences in baseline mite levels). No colonies were treated between study commencement and termination, and no colonies were treated with Fumagillin-B (a product for controlling *Vairimorpha* spp.) or oxytetracycline (an antibiotic for controlling EFB disease).

The *Varroa* validation dataset consisted of previously published data quantifying *Varroa* infestations in Canada in 2016 (described by Borba *et al.*^47^). We used these data to validate our findings relating *Varroa* to regional, temporal, and climate variables and explore how length of beekeeping season relates to *Varroa* occurrence. For our analysis, we only used *Varroa* infestation data (mites per 100 bees, determined by the alcohol wash method^47^) from overwintered colonies belonging to, and managed by, beekeeper collaborators where their mite loads were not manipulated. This includes N = 480 overwintered colonies located in BC, southern AB, southern ON, and southern QC. Colonies within BC were located in 11 yards and were owned by 11 operators in coastal BC (<100 km from the coast; 8 yards) and interior BC (Thompson/Okanagan and Kootenay/Boundary regions; 3 yards), which have notably different climates. Colonies in AB, ON, and QC were also distributed across multiple yards, all of which are shown in Borba *et al.*^47^ Each colony was sampled at up to three time points during the active beekeeping season. Data were filtered to exclude colonies from which fewer than two mite wash samples were obtained between May and September, as well as colonies which beekeepers moved to unknown locations between mite sampling events.

### Pathogen testing (primary dataset)

*Varroa, M. plutonius,* and *Vairimorpha* 2020 and 2021 sample analysis was performed exactly as previously described.^44^ Briefly, for *M. plutonius* and *Vairimorpha*, samples were analyzed as pooled replicates (15 bees from 4 colonies each, yielding n = 5 replicates from each site type (near to or far from focal crops) for each crop in each year. The same four colonies were pooled at each time point, and pooled colonies were always located together in the same yard (and experienced the same climatic conditions). *M. plutonius* analysis was performed by endpoint PCR and *Vairimorpha* spore counts were determined by microscopy (hemocytometer counting) and expressed as number of spores per bee.^44^ Samples for *Varroa* mite counts were not pooled across colonies, and mite counts per 100 bees were determined using the alcohol wash method as previously described.^47^ Therefore, final sample sizes for *M. plutonius* and *Vairimorpha* data were N = 60 pooled samples per time point, per year (360 samples from 120 pooled four-colony units), whereas the sample size for *Varroa* was N = 240 colonies per time point, per year (1,440 samples from 480 colonies). Please see the previously published methods for specific details regarding the protocols used.^44^

### Climatic data extraction

Datasets from Environment and Climate Change Canada climate stations^48^ were used to extract hourly and daily climate variables (mean, maximum, and minimum daily temperatures, average daytime and nighttime wind speeds, and total precipitation). The climate stations that were within 30 km of the colony locations were selected and the daily and hourly datasets for the corresponding time of the sampling events were downloaded. The daily climate records were used to extract the daily climate variables for the sampling dates and were averaged over three weeks (21 days) prior to the sampling date. Averaging climatic variables across time prior to sampling dates improves model fits,^9^ and three weeks prior (corresponding to one honey bee brood cycle) was the longest period for which we had precise knowledge of colony GPS locations. Because wind is predominantly expected to influence honey bees via changing their flight behavior, and since flight only occurs during the daylight hours, we used the hourly climate records to extract daytime and nighttime averages of wind speed for the sampling dates, which were also averaged over three weeks prior to the sampling dates. Daytime was defined as 5 am to 8 pm for all locations and dates, except for September sampling in Ontario, for which daytime was defined as 6 am to 7 pm to more closely match actual sunrise and sunset times late in the season. For the colonies with more than one climate station within a 30 km distance, the daily, daytime, and nighttime climate data for sampling dates and three weeks prior to the sampling dates were extracted for each climate station independently, then spatially averaged across all climate stations within the 30 km distance.

Prior to conducting statistical modelling, correlations among climate variables in the primary dataset were assessed by computing Pearson correlation coefficients (**Supplementary Figure S2**). Based on these results, we included only mean temperature, average daytime wind speed, and total precipitation in our statistical models (described under “Statistical analysis,” as these variables had correlation coefficients of |r| < 0.4 in 2020 and < 0.3 in 2021. We chose to use daytime wind exclusively, because wind is expected to only impact honey bees during the day when they have an opportunity to fly. For precipitation, we used daily totals, rather than daytime totals, because precipitation is also linked to nectar production in flowers and ambient humidity regardless of when it falls, and could therefore conceivably affect honey bee behavior, thermoregulation, or nutrition at any time.

### Statistical analysis

All statistical analyses were performed in R (version 4.3.0) using R Studio (version 2023.09.1+494)^49^. The primary dataset (2020 and 2021) and *Varroa* validation dataset (2016) were analyzed separately. For the primary dataset, site type (whether colonies/replicates were located near or far from focal crops) was first considered as a covariate but in no case was it a significant predictor, so it was dropped from the models. For all models, appropriateness of fit was confirmed using tools within the DHARMa package^50^ Interaction terms were removed from the model if they were non-influential and if appropriateness of fit was retained after doing so.

The *Varroa* data in the primary dataset are proportional (mites per 100 bees) and zero-heavy; therefore, we analyzed the data in two ways: first, we modelled the non-zero subset of the data (153/480 samples) to describe how intensity of *Varroa* infestation changed in different regions over time, then, we modelled binomial *Varroa* detections (occurrence) separately. In no cases (for *Varroa* nor other pathogens) were zeroes imputed. The non-zero *Varroa* data were transformed by taking the natural logarithm and analyzed using a linear mixed model (lme4 package^51^) with year (two levels) as a fixed effect, calendar date (continuous) and general region/operation (eight levels) as interacting effects, and colony as a random intercept term. Next, *Varroa* occurrence (1 = detected, 0 = not detected) from sampling dates at similar points in the beekeeping season (all in July for 2020 and all in June for 2021, no repeated sampling, years analyzed separately) were extracted from the complete dataset and analyzed using a generalized linear model, specifying a binomial distribution, and with region/operation (five levels in each year) included as a fixed effect. The Ontario and Quebec-Montreal regions were excluded from this analysis as they did not have a *Varroa* sample taken at a comparable date to the other regions.

To determine relationships with regional, temporal and climatic variables in the primary dataset, *Varroa* occurrence from all time points and regions were analyzed using a generalized linear mixed model according to the following formula: Occurrence ∼ Temperature*Wind+Precipitation + Date*Region + Year + (1|Colony). Here and in all cases (except for the *Varroa* validation analysis; see below), the three climatic variables were initially included as a three-way interaction term, but the model was simplified if this interaction was not significant. If a climate variable did not significantly predict occurrence as a main or interactive effect, and if retaining the variable in the model did not significantly improve residual or simulated residual distributions, the variable was dropped. *M. plutonius* occurrence was modelled using the same principles, according to the formula: Occurrence ∼ Wind*Precipitation + Date*Region + Year + (1|Unit), where unit refers to the pooled sample group. For *Vairimorpha*, non-zero intensities (spore loads; 305/360 samples) were first log10 transformed, then modelled using a linear mixed model according to the formula: Intensity ∼ Wind*Precipitation + Temp + Date*Region + Year + (1|Unit). *Vairimorpha* data from Quebec in 2020 were removed as there were too few data points to draw meaningful comparisons (< n=3 per site type and time point).

To validate the results of our primary analysis of *Varroa* occurrence and climate data, we used the same linear model to describe *Varroa* occurrence data on the 2016 validation dataset (except that year was not included, as this dataset only spans one year). This dataset comprised *Varroa* occurrence previously published by Borba *et al.*^47^ (see “honey bee colonies and datasets” section). The climate data were downloaded and averaged for each site in the same manner as described for the 2020 and 2021 data. In a second evaluation, we analyzed *Varroa* occurrence at the three different sampling events (early season, mid-season, and late season) separately, as the mid-season sampling dates have the greatest temporal overlap with the 2020 and 2021 data. For this analysis, we used a generalized linear model (no random effects were specified as there was no repeated sampling of colonies within each sampling event). Mean daily temperature data (spatially averaged across climate stations within a 30 km radius of each site) for full 2016 calendar year were also obtained for the corresponding GPS locations and used to estimate the beginning and end of the brood rearing season. While we do not know the actual dates that substantial brood rearing commenced and ceased, we estimated it to begin on the date at which the seven-day sliding average temperature was at or above 5°C (when brood rearing intensifies^52^) for at least three consecutive days, and ending on the date at which the seven-day sliding average temperature declined below 5°C for ten or more consecutive days. We chose these criteria on the rationale that it guards against making inferences based on spurious fluctuations. While imprecise, the approach is meant to illustrate broad patterns of seasonal timing in different regions.

### Data availability

All raw data underlying these analyses are available as supplementary material. 2020 and 2021 pathogen, parasite, regional, temporal, climate, and GPS location data are available in **Supplementary Data 1 & 2**, respectively. Colony locations are also depicted in **Supplementary Files 1 & 2.** The 2016 *Varroa* and climate data are reported in **Supplementary Data 3**.

## Results

Analysis of the primary dataset shows that *Varroa* intensity generally increased over time, as expected, but varied significantly by region and year (**Figure 1A**; see **Table 1** for statistical reporting). Notably, *Varroa* intensities in BC’s Lower Mainland were generally high even early in the season (April-May), with levels comparable to late-season (Aug-Sept) intensities in other provinces. Indeed, selecting comparable sampling dates across provinces (July sampling in 2020 and June sampling in 2021) shows that *Varroa* prevalence in BC is high with respect to other provinces, with an 88% and 50% positivity rate in 2020 and 2021, respectively (**Figure 1B & C**).

**Figure 1.**
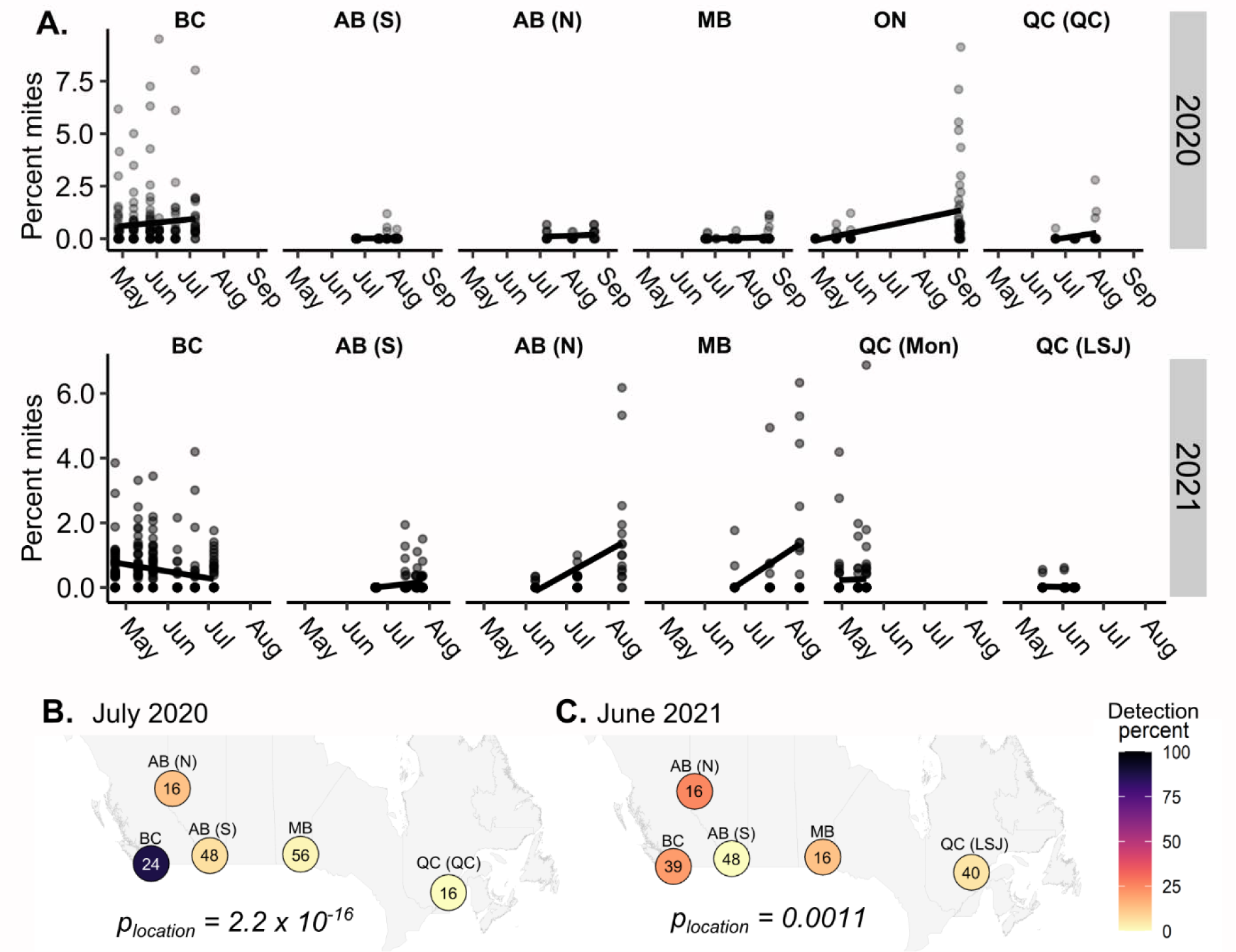
Regional patterns of *Varroa destructor* intensity and occurrence. A) *Varroa* intensities (non-zero mites per 100 bees) varied significantly by year and a date-by-region interaction. Statistical parameters are described in **Table 1**. BC = British Columbia, AB (S) = Southern Alberta, AB (N) = Northern Alberta, MB = Manitoba, ON = Ontario, QC (QC) = Quebec, near Quebec City, QC (Mon) = Quebec, near Montreal, QC (LSJ) = Quebec, near Lac St. Jean. B & C) Region effects illustrated as occurrence rates at each site, restricted to comparable sampling dates (July dates in 2020 and June dates in 2021, no repeated sampling). Logistic regression analysis shows significant regional differences in occurrence frequencies in 2020 (C; χ^2^ = 82.6, df = 4, p < 0.001) and less pronounced, but still significant differences in 2021 (D; χ^2^ = 18.3, df = 4, p = 0.0011). Numbers inside circles indicate sample sizes.

**Table 1.** Linear mixed model of non-zero mite intensities^a^ by date, region, and year.

| **Analysis of deviance table (Type II Wald $\chi^2$)** | **$\chi^2$** | **df** | **p** | **Code<sup>b</sup>** |
| --- | --- | --- | --- | --- |
| Date<sup>c</sup> | 15.8 | 1 | < 0.001 | *** |
| Region | 25.7 | 7 | < 0.001 | *** |
| Year | 5.0 | 1 | 0.026 | * |
| Date x Region | 22.7 | 7 | 0.0019 | ** |
| **Parameter** | **Coef** | **t<sub>(366)</sub>** | **p** | **Code** |
| (Intercept) [Date=0, Region=BC, Year=2020] | -0.39 | -4.04 | < 0.001 | *** |
| Date | 0.06 | 0.75 | 0.45 |  |
| Region [AB S] | 0.66 | 0.78 | 0.43 |  |
| Region [AB N] | -0.99 | -4.33 | < 0.001 | *** |
| Region [MB] | -0.86 | -2.35 | 0.019 | * |
| Region [ON] | -0.06 | -0.26 | 0.80 |  |
| Region [QC QC] | -0.39 | -0.46 | 0.65 |  |
| Region [QC Mon] | -0.1 | -0.16 | 0.87 |  |
| Region [QC LSJ] | -0.26 | -0.21 | 0.84 |  |
| Year [2021] | 0.24 | 2.23 | 0.026 | * |
| Date x Region [AB S] | -1.27 | -1.44 | 0.15 |  |
| Date x Region [AB N] | 0.57 | 3.47 | < 0.001 | *** |
| Date x Region [MB] | 0.81 | 3.05 | 0.0020 | ** |
| Date x Region [ON] | 0.16 | 1.20 | 0.23 |  |
| Date x Region [QC QC] | 1.02 | 1.18 | 0.24 |  |
| Date x Region [QC Mon] | -0.09 | -0.17 | 0.86 |  |
| Date x Region [QC LSJ] | 0.23 | 0.13 | 0.89 |  |
| **Random effect** | **Variance** | **sd** | **Levels** | **Obs** |
| Random intercept [Colony] | 0.31 | 0.55 | 232 | 385 |
| *Full model $R^2 = 0.57$* |  |  |  |  |
| *Formula = $\ln(\text{mite percent}) \sim \text{Year} + \text{Date} * \text{Region} + (1 \vert \text{Colony})$* |  |  |  |  |
| *Family = Gaussian* |  |  |  |  |

<sup>a</sup>Mite intensities (non-zero mites per 100 bees) were transformed by taking the natural logarithm696 <sup>b</sup>Significance codes: p < 0.001 = \*\*\*, p < 0.01 = \*\*, p < 0.05 = \*697 <sup>c</sup>Date was scaled prior to modelling

We next modelled *Varroa* occurrence with regional, temporal (date, and year) and climatic (mean temperature, average wind speed, and total precipitation) predictors to determine how weather patterns influence occurrence probability. Date and temperature were, predictably, strongly correlated with one another (Pearson’s r = 0.9; **Supplementary Figure S2**). Nevertheless, temperature still positively predicted *Varroa* occurrence beyond the covarying effect of date. Interestingly, although *Varroa* occurrence was generally high in BC’s Lower Mainland, occurrence probability did not increase appreciably with time, as in most other regions (**Figure 2 A & B**). We also found a significant interactive effect between temperature and wind, with higher wind speeds predicting lower *Varroa* occurrence probability, but this effect was moderated by high temperatures (**Supplementary Figure S3**). Precipitation was also negatively associated with *Varroa* occurrence, and the lowest occurrence probabilities were observed when both wind speeds and precipitation were high (**Figure 2C**). This effect was consistent across 2020 and 2021. See **Table 2** for complete statistical reporting.

**Figure 2.**
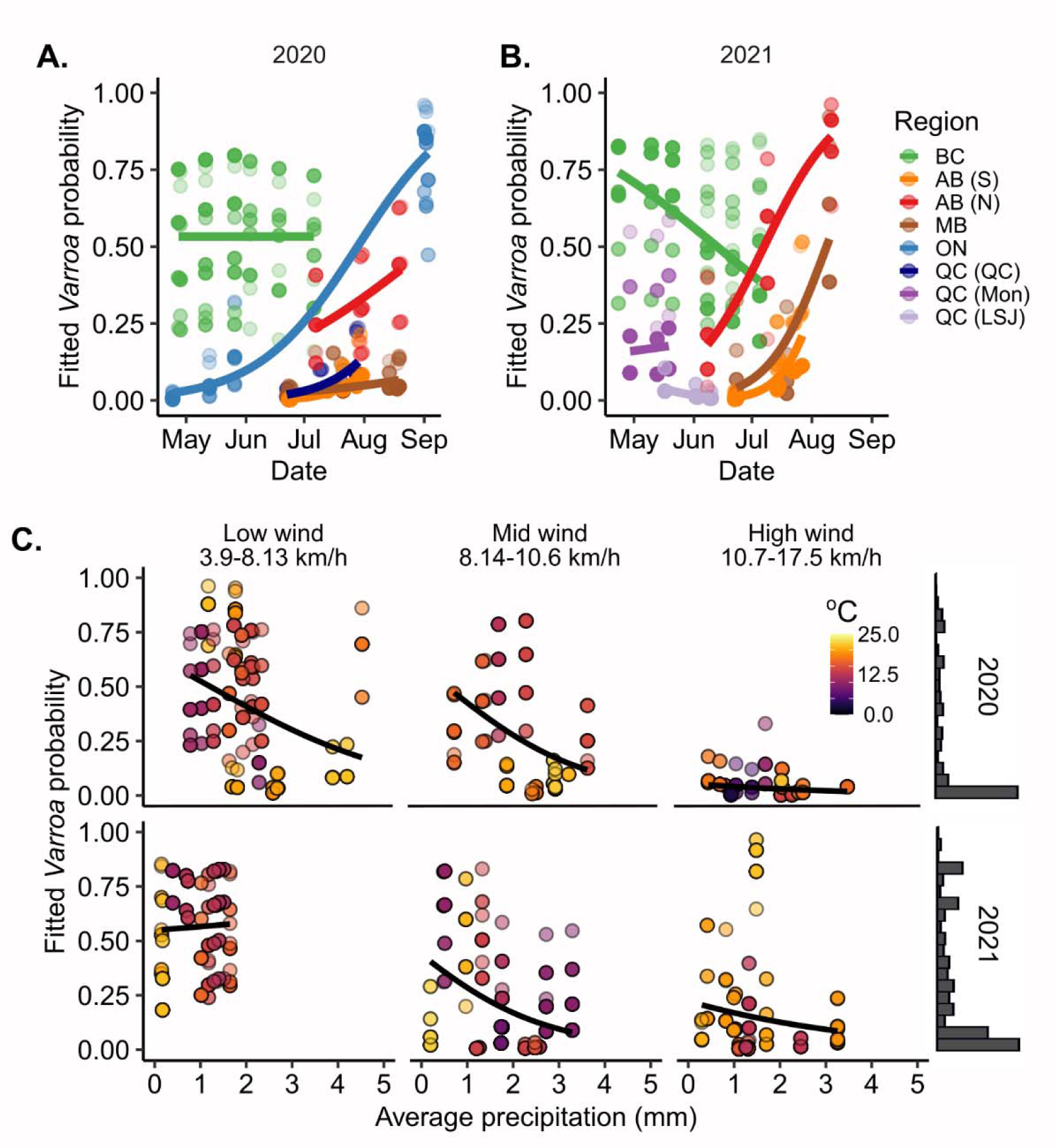
*Varroa destructor* occurrence is linked to regional, temporal and climatic variables. A & B) Fitted *Varroa* occurrence probabilities (1 = present, 0 = absent) vary by year and a date-by-region interaction, among other influential factors. Statistical parameters fully reported in **Table 2**. C) Occurrence probabilities significantly decrease with higher wind speeds, but the effect is reduced when temperatures are warm (see also **Supplementary Figure S2**). Occurrence also significantly decreases with precipitation. Histograms show densities of fitted probabilities.

**Table 2.** Generalized linear mixed model of mite occurrence.

| **Analysis of deviance table (Type II Wald $\chi^2$)** | **$\chi^2$** | **df** | **p** | **Code<sup>a</sup>** |
| --- | --- | --- | --- | --- |
| Temperature<sup>b</sup> | 4.3 | 1 | 0.037 | * |
| Wind<sup>b</sup> | 3.0 | 1 | 0.084 |  |
| Precipitation<sup>b</sup> | 9.2 | 1 | 0.0024 | ** |
| Date<sup>b</sup> | 6.1 | 1 | 0.013 | * |
| Region | 85.7 | 7 | < 0.001 | *** |
| Year | 6.5 | 1 | 0.011 | * |
| Temperature x Wind | 4.8 | 1 | 0.029 | * |
| Date x Region | 29.6 | 7 | < 0.001 | *** |
| **Parameter** | **Coef** | **z** | **p** | **Code** |
| (Intercept) [Temperature=0, Wind=0, Precipitation=0, Date=0, Region=BC, Year=2020] | 0.37 | 1.0 | 0.31 |  |
| Temperature | 0.73 | 2.3 | 0.023 | * |
| Wind | 0.61 | 2.3 | 0.022 | * |
| Precipitation | 0.27 | 3.0 | 0.0024 | ** |
| Date | -0.44 | -1.2 | 0.23 |  |
| Region [AB S] | -8.00 | -5.9 | < 0.001 | *** |
| Region [AB N] | -2.23 | -3.5 | < 0.001 | *** |
| Region [MB] | -5.40 | -6.0 | < 0.001 | *** |
| Region [ON] | -0.81 | -1.3 | 0.18 |  |
| Region [QC QC] | -3.86 | -2.6 | 0.0095 | ** |
| Region [QC Mon] | -2.12 | -1.6 | 0.11 |  |
| Region [QC LSJ] | -6.95 | -4.5 | < 0.001 | *** |
| Year [2021] | 0.67 | 2.5 | 0.011 | * |
| Temperature x Wind | 0.47 | 2.2 | 0.029 | * |
| Date x Region [AB S] | 4.10 | 3.7 | < 0.001 | *** |
| Date x Region [AB N] | 1.39 | 2.7 | 0.0077 | * |
| Date x Region [MB] | 1.53 | 2.6 | 0.011 | * |
| Date x Region [ON] | 1.40 | 3.9 | < 0.001 | *** |
| Date x Region [QC QC] | 2.01 | 1.3 | 0.20 |  |
| Date x Region [QC Mon] | 0.48 | 0.4 | 0.67 |  |
| Date x Region [QC LSJ] | -3.52 | -1.8 | 0.079 |  |
| **Random effect** | **Variance** | **sd** | **Levels** | **Obs** |
| Random intercept [Colony] | 1.56 | 1.25 | 479 | 1426 |
| *Full model $R^2 = 0.64$* |  |  |  |  |
| *Formula = Occurrence ~ Temperature\*Wind+Precipitation + Date\*Region + Year + (1\|Colony)* |  |  |  |  |
| *Family = Binomial* |  |  |  |  |

<sup>a</sup>Significance codes: p < 0.001 = \*\*\*, p < 0.01 = \*\*, p < 0.05 = \*701 <sup>b</sup>Variables were scaled (z scored) prior to fitting

To determine if our findings agreed with an independent dataset, we analyzed a validation dataset of *Varroa* occurrence acquired in 2016 from BC, AB, ON, and QC using the same model parameters. Unlike the 2020 and 2021 data, this dataset covers a longer time period, spanning dates from April to November (**Figure 3A**). Climatic variables in the primary and validation datasets had comparable average magnitudes, although the maximum precipitation observed in the primary dataset was notably higher than the validation dataset (primary dataset: mean temperature = 15.1 °C (2.9-22.7 °C), mean precipitation = 2.0 mm (0.1-19.6 mm), mean wind speed = 9.5 km/h (3.9-17.5 km/h); validation dataset: mean temperature = 15.6 °C (6.7-23.0 °C), mean precipitation = 2.4 mm (0.1-9.5 mm), mean wind speed = 11.6 km/h (3.4-21.3 km/h)). Early season (April-May) occurrence probabilities were again highest in BC (coastal regions including the Lower Mainland) and low in QC, which might reflect differences in brooding initiation in these areas, as temperatures in BC yards exceeded 5°C earlier than yards in all other regions (**Supplementary Figure S4**); however, wind was not associated with an overall decrease in occurrence (**Supplementary Figure S5**). Rather, temperature was the ultimate climatic predictor (associated with higher occurrence probability), again above and beyond a simple covariation between temperature and date (see **Table 3** for complete statistical reporting). We hypothesized that this may be due to the dataset covering a wider range of temperatures across a longer season. Indeed, when we inspected early, mid, and late-season occurrence separately, the mid-season occurrence – which overlap with the 2020 and 2021 sampling dates to the greatest degree – do display a negative relationship with wind, but no significant relationship with precipitation (**Figure 3B**).

**Figure 3.**
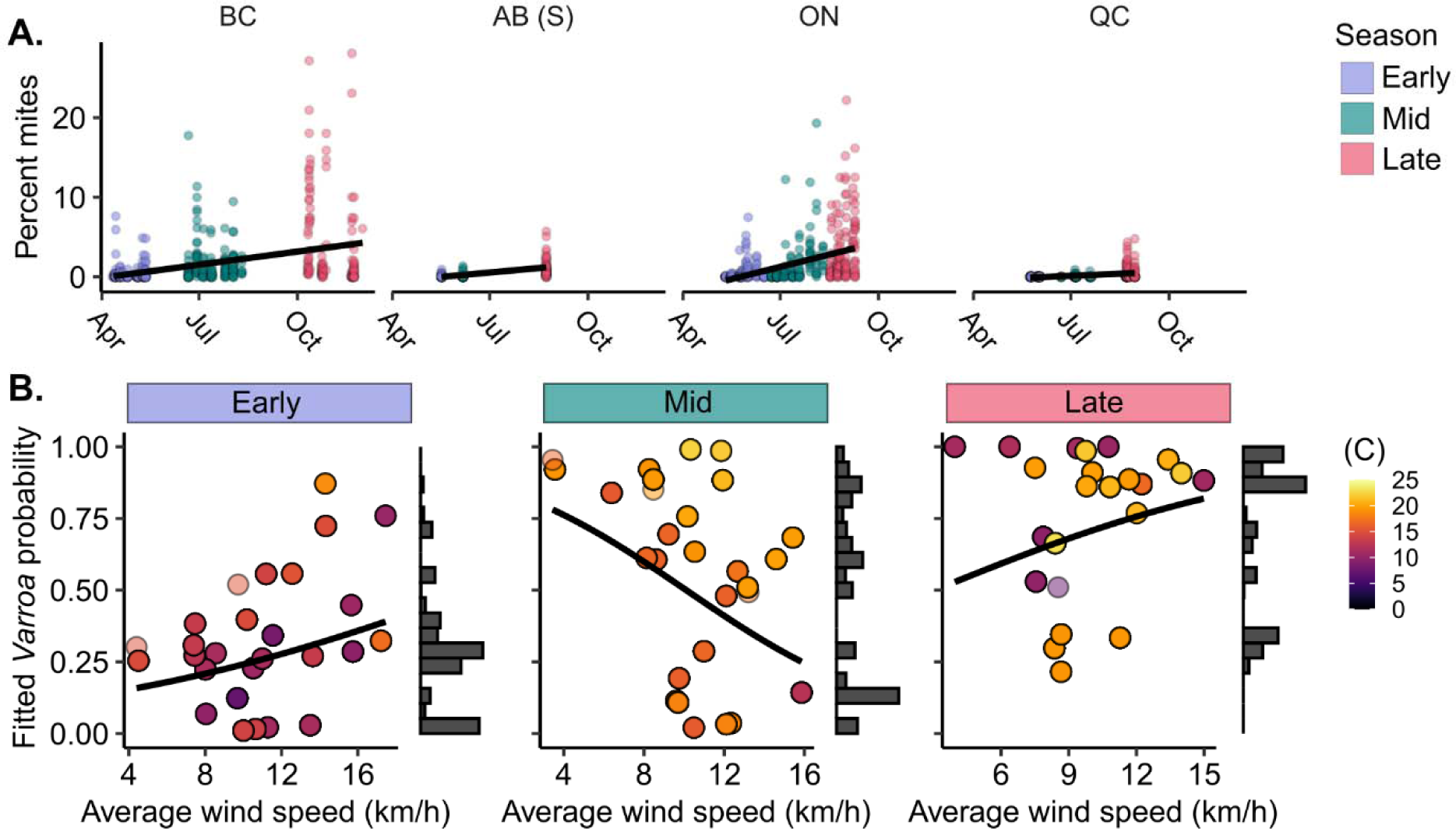
Validation analysis: *Varroa* occurrence probability decreases with wind mid-season, but temperature drives overall trends. A) Validation dataset of *Varroa* intensity and occurrence in BC, AB, ON, and QC during the 2016 beekeeping season (divided by early, mid-season, and late sampling events). Data were modelled according to the same formula as the 2020/2021 dataset, with results described in **Table 3**. The positive effect of temperature on *Varroa* occurrence probability was exacerbated, not mitigated, under windier conditions (**Supplementary Figure S4**). B) Examining each sampling time point separately, only mid-season *Varroa* occurrence (n = 480) is negatively associated with wind. Mid-season sampling (June to mid-August) dates have the greatest overlap with 2020 and 2021 sampling dates. Model parameters and outcomes for mid-season data are reported in **Table 4**.

**Table 3.**
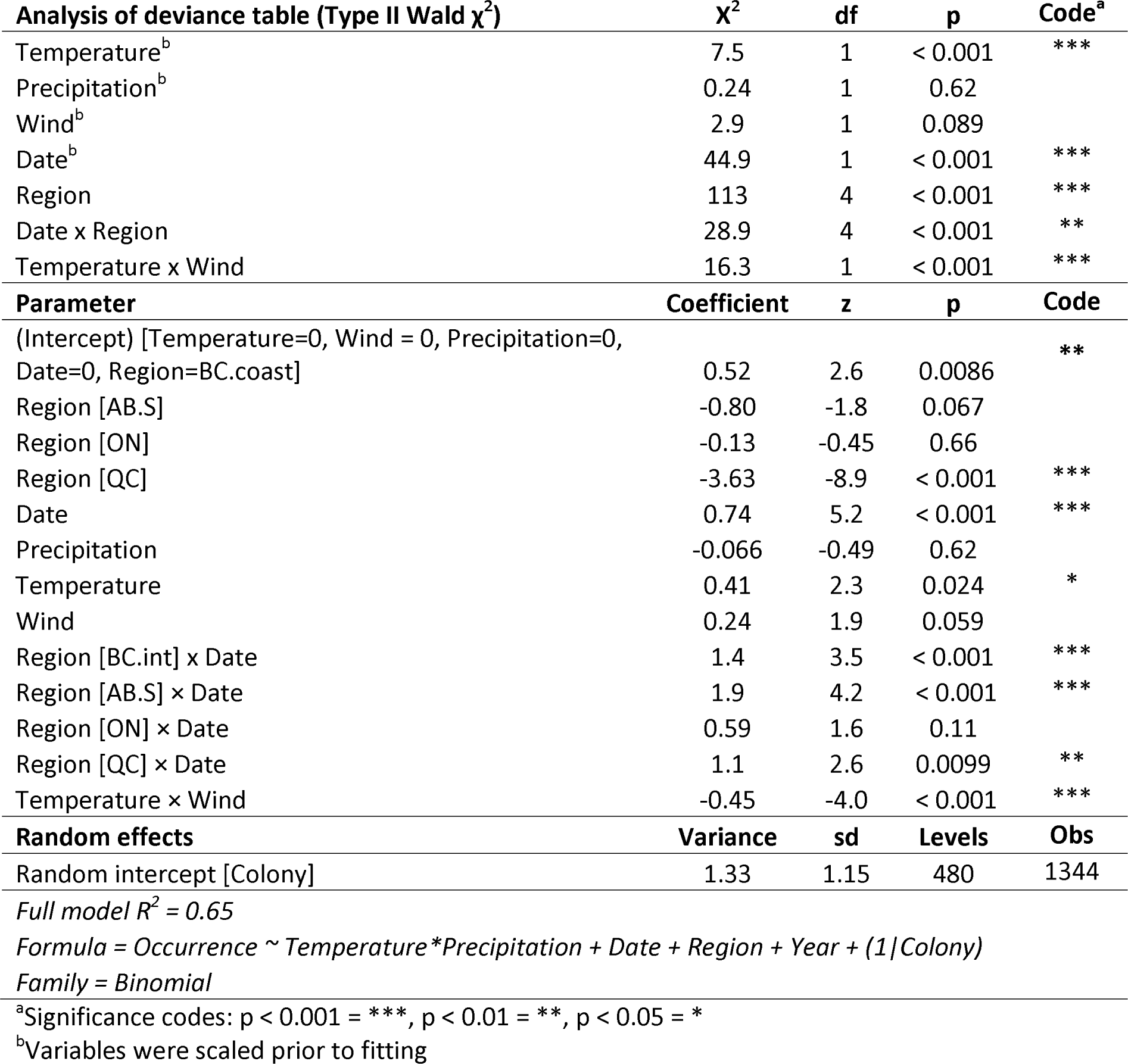
*Varroa* model validation: Full-season data.

**Table 4.**
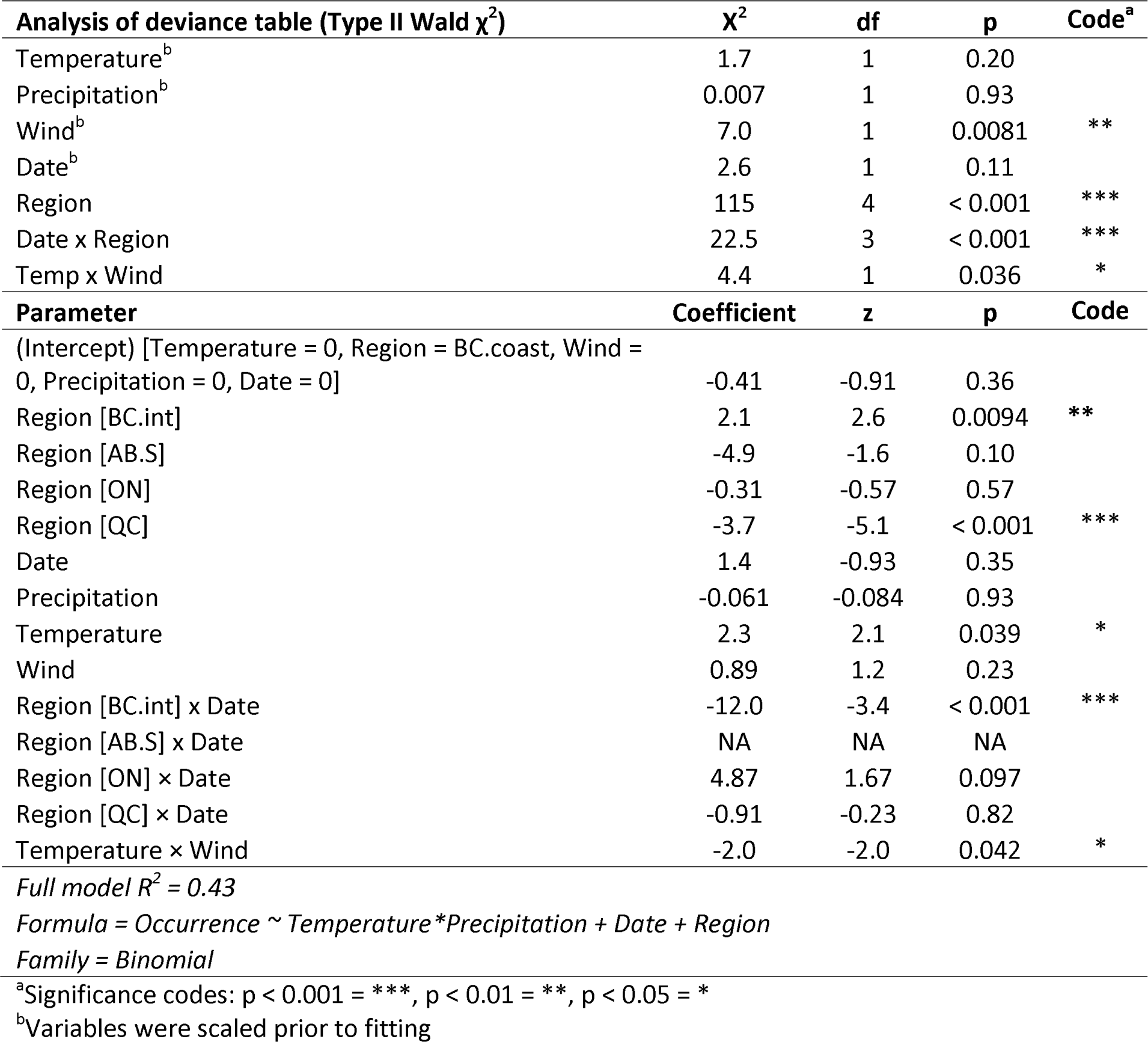
*Varroa* model validation: Mid-season data.

Next, we used the primary dataset to analyze relationships between *Vairimorpha* and the same regional, temporal and climatic variables as for *Varroa*. Spores were detected in 95% of replicates (114/120) for at least one time point; therefore, we used a linear mixed model with regional, temporal, and climatic variables as predictors of intensity and did not model *Vairimorpha* presence/absence. We again observed strong regional effects, but the patterns varied greatly from year to year (**Figure 4A & B**; see **Table 5** for statistical reporting). For example, while patterns were similar in BC and southern AB across years, samples from northern AB and MB tended to have higher intensities in 2020 than 2021. While patterns over time within regions varied, overall spore intensities decreased as temperatures warmed and the season progressed (**Figure 4C**), but the effect of temperature was marginally not significant (Type II Wald χ^2^ test; χ^2^ = 3.09, df = 1, N = 480, p = 0.079) after accounting for the effect of date (**Table 5**). Wind speed and precipitation significantly interacted, with spore intensities increasing with higher precipitation, but only under windier conditions (**Figure 4D**).

**Figure 4.**
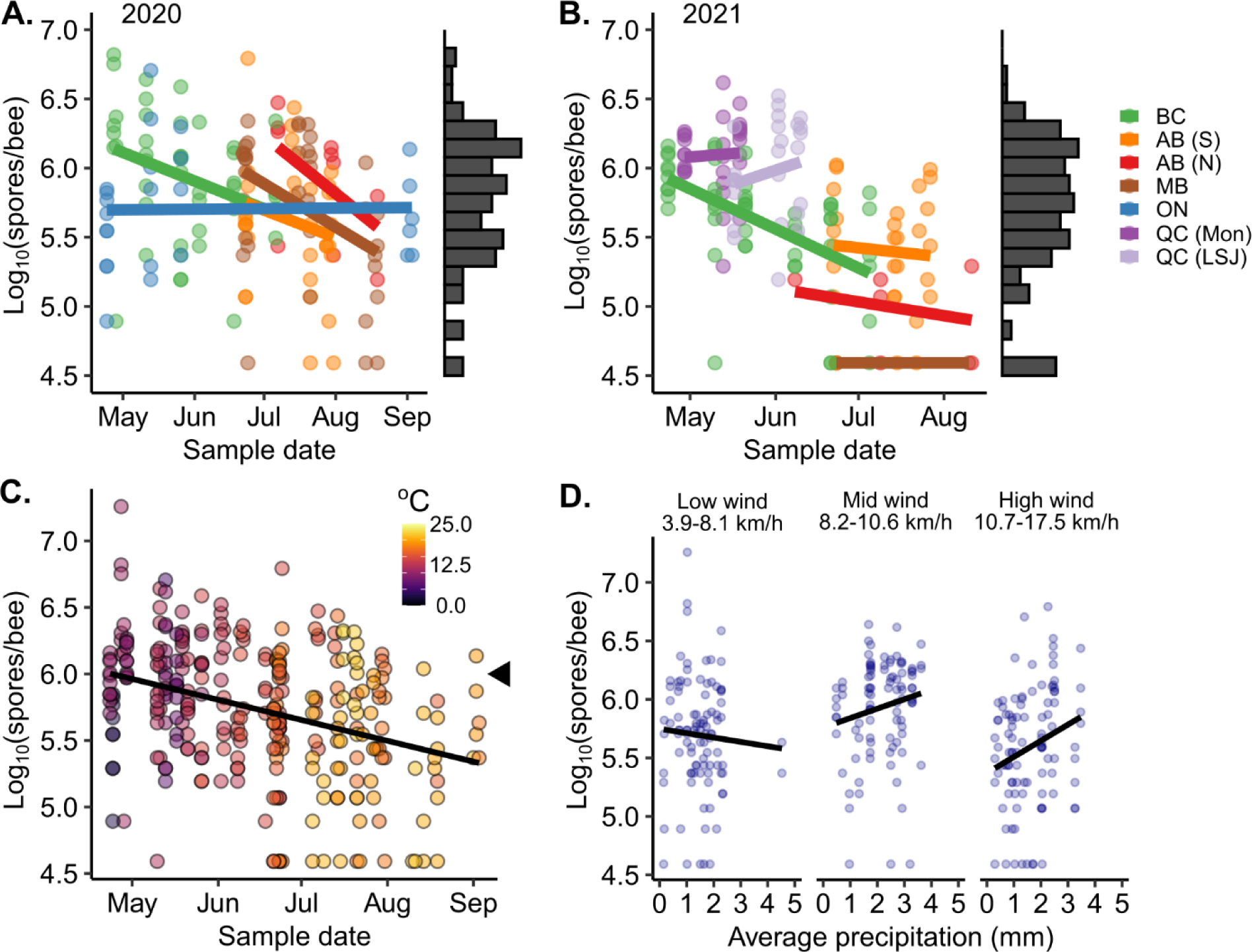
*Vairimorpha* spore intensities are linked to regional, temporal and climatic variables. Statistical parameters fully reported in **Table 3**. Spore intensities vary significantly by region in 2020 (A) and 2021 (B). Spore intensities generally decrease over time during the spring and summer, but this trend varies by region. Histograms show spore intensity bins. C) The general decrease over time coincides with increased temperatures. The black triangle indicates the economic threshold of 1 million spores/bee. D) Spore intensities significantly increase with precipitation, but only under windy conditions, illustrated in three equidensity wind speed bins. BC = British Columbia, AB (S) = Southern Alberta, AB (N) = Northern Alberta, MB = Manitoba, ON = Ontario, QC (Mon) = Quebec, near Montreal, QC (LSJ) = Quebec, near Lac St. Jean.

**Table 5.** Linear mixed model of *Vairimorpha* spp. spore intensities^a.^

| **Analysis of deviance table (Type II Wald $\chi^2$)** |  |  |  |  |
| --- | --- | --- | --- | --- |
|  | **$\chi^2$** | **df** | **p** | **Code<sup>b</sup>** |
| Wind<sup>c</sup> | 0.00 | 1 | 0.99 |  |
| Precipitation<sup>c</sup> | 0.53 | 1 | 0.47 |  |
| Temperature<sup>c</sup> | 3.09 | 1 | 0.079 |  |
| Date<sup>c</sup> | 21.4 | 1 | < 0.001 | *** |
| Region | 38.5 | 6 | < 0.001 | *** |
| Year | 32.3 | 1 | < 0.001 | *** |
| Wind x Precipitation | 9.41 | 1 | 0.0020 | ** |
| Date x Region | 14.2 | 6 | 0.027 | * |
| **Parameter** | **Coefficient** | **t** | **p** | **Code** |
| (Intercept) [Wind= 0, Precipitation=0, Temperature=0, Date=0, Region=BC, Year=2020] | 5.7 | 54.7 | < 0.001 | *** |
| Wind | -0.02 | -0.31 | 0.76 |  |
| Precipitation | 0.02 | 0.79 | 0.43 |  |
| Temperature | 0.17 | 1.76 | 0.080 |  |
| Date | -0.46 | -4.37 | < 0.001 | *** |
| Region [MB] | 0.06 | 0.35 | 0.73 |  |
| Region [AB N] | 0.16 | 0.70 | 0.49 |  |
| Region [ON] | 0.26 | 1.49 | 0.14 |  |
| Region [QC LSJ] | 0.74 | 3.41 | < 0.001 | ** |
| Region [QC Mon] | 0.92 | 2.62 | 0.009 | * |
| Region [AB S] | 0.18 | 0.66 | 0.51 |  |
| Year [2021] | -0.45 | -5.68 | < 0.001 | *** |
| Wind x Precipitation | 0.10 | 3.07 | 0.002 | ** |
| Scaled x Region [MB] | 0.06 | 0.49 | 0.62 |  |
| Scaled x Region [AB N] | 0.19 | 0.96 | 0.34 |  |
| Scaled x Region [ON] | 0.31 | 3.25 | 0.001 | ** |
| Scaled x Region [QC LSJ] | 0.29 | 0.99 | 0.32 |  |
| Scaled x Region [QC Mon] | 0.37 | 1.25 | 0.21 |  |
| Scaled x Region [AB S] | 0.11 | 0.65 | 0.52 |  |
| **Random effect** | **Variance** | **sd** | **Levels** | **Obs** |
| Random intercept [Unit] | 0.043 | 0.21 | 114 | 296 |
| *Full model $R^2 = 0.50$* |  |  |  |  |
| *Formula = Spores ~ Wind\*Precipitation + Temperature + Date\*Region + Year + (1\|Unit)* |  |  |  |  |
| *Family = Gaussian* |  |  |  |  |

<sup>a</sup>Spore intensities were log<sub>10</sub> transformed
<sup>b</sup>Significance codes: p < 0.001 = \*\*\*, p < 0.01 = \*\*, p < 0.05 = \*
<sup>c</sup>Variables were scaled prior to fitting

*M. plutonius* occurrence varied by region as well, but there was no significant effect of year (**Table 6**). Overall detection probability was high in BC and markedly increased over time between May and June, whereas zero detections were recorded in ON throughout the season (**Figure 5A**). Temperature was not a significant predictor after accounting for date, and was removed from the model; however, wind and precipitation significantly interacted, with increased *M. plutonius* occurrence probability under wetter, windier conditions as well as drier, less windy conditions, and lower occurrence probability under intermediate conditions **(Figure 5B**).

**Figure 5.**
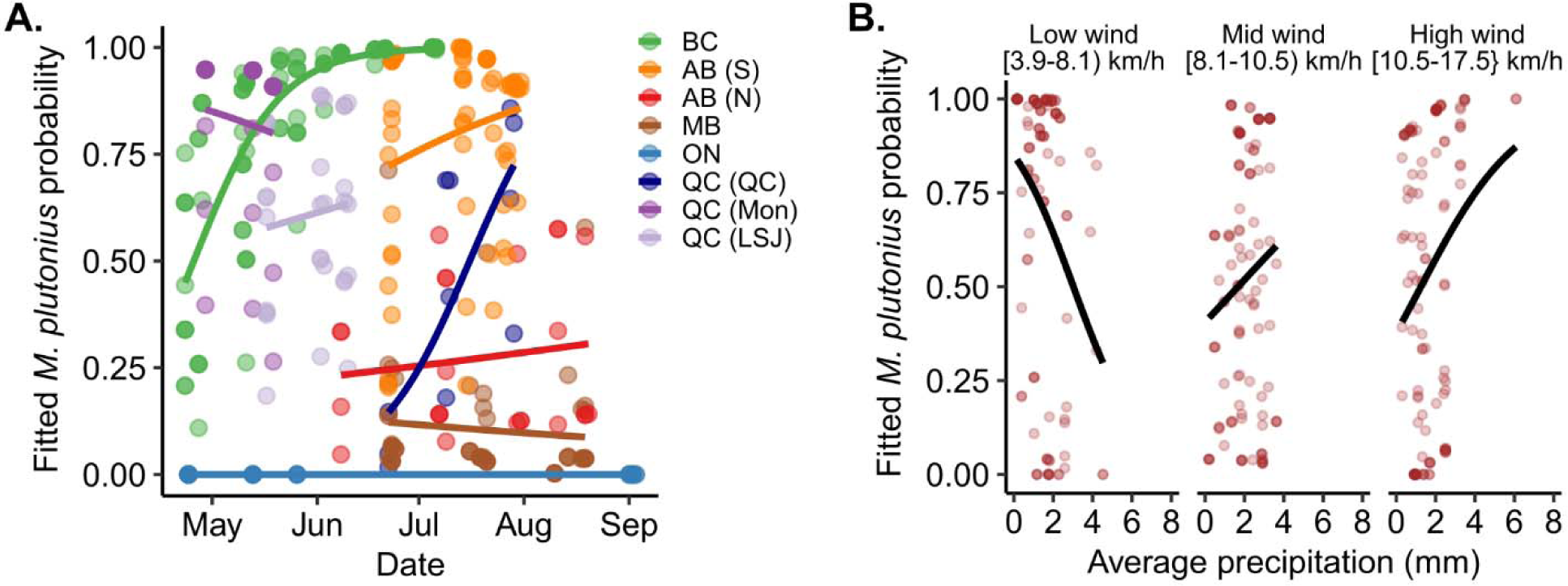
*Melissococcus plutonius* occurrence is linked to region, precipitation, and temperature. Occurrence was fitted with generalized linear mixed model with statistical parameters reported in **Table 4**. A) Occurrence varies significantly by region. B) Wind and precipitation significantly interact, as illustrated by plotting fitted probabilities against precipitation in three equidensity wind bins. *M. plutonius* occurrence decreases with precipitation when wind speed is low, but increase with precipitation when wind is medium-high. BC = British Columbia, AB (S) = Southern Alberta, AB (N) = Northern Alberta, MB = Manitoba, ON = Ontario, QC (QC) = Quebec, near Quebec City, QC (Mon) = Quebec, near Montreal, QC (LSJ) = Quebec, near Lac St. Jean.

**Table 6.** Generalized linear mixed model of *M. plutonius* occurrence.

| **Analysis of deviance table (Type II Wald $\chi^2$)** |  |  |  |  |
| --- | --- | --- | --- | --- |
|  | **$\chi^2$** | **df** | **p** | **Code<sup>a</sup>** |
| Wind<sup>b</sup> | 0.0001 | 1 | 0.99 |  |
| Precipitation<sup>b</sup> | 0.41 | 1 | 0.52 |  |
| Date<sup>b</sup> | 5.4 | 1 | 0.021 | * |
| Region | 25.0 | 7 | < 0.001 | *** |
| Year | 0.75 | 1 | 0.39 |  |
| Wind x Precipitation | 6.7 | 1 | 0.0099 | ** |
| Date x Region | 12.1 | 7 | 0.096 |  |
| **Parameter** | **Coefficient** | **z** | **p** | **Code** |
| (Intercept) [Wind = 0, Precipitation=0, Date=0, Region=BC] | 5.8 | 4.4 | < 0.001 | *** |
| Wind | 0.53 | 0.72 | 0.47 |  |
| Precipitation | 0.21 | 0.41 | 0.68 |  |
| Date | 3.8 | 3.8 | < 0.001 | *** |
| Region [MB] | -8.6 | -4.8 | < 0.001 | *** |
| Region [AB.N] | -7.3 | -4.0 | < 0.001 | *** |
| Region [ON] | -24.7 | -0.03 | 0.97 |  |
| Region [QC.QC] | -7.1 | -3.1 | 0.002 | ** |
| Region [QC.LSJ] | -4.7 | -2.5 | 0.012 | * |
| Region [QC.Mon] | -4.6 | -1.5 | 0.14 |  |
| Region [AB.S] | -4.5 | -1.7 | 0.084 |  |
| Year [2021] | -0.56 | -0.86 | 0.39 |  |
| Wind x Precipitation | 2.2 | 2.6 | 0.01 | * |
| Date x Region [MB] | -3.7 | -2.9 | 0.004 | ** |
| Date x Region [AB.N] | -3.3 | -2.6 | 0.011 | * |
| Date x Region [ON] | -3.8 | -0.0051 | 1.0 |  |
| Date x Region [QC.QC] | 1.2 | 0.53 | 0.60 |  |
| Date x Region [QC.LSJ] | -3.7 | -1.9 | 0.056 |  |
| Date x Region [QC.Mon] | -5.2 | -1.9 | 0.06 |  |
| Date x Region [AB.S] | -2.5 | -1.5 | 0.15 |  |
| **Random effects** | **Variance** | **sd** | **Levels** | **Obs** |
| Random intercept [Unit] | 2.7 | 1.6 | 120 | 360 |
| *Full model $R^2 = 0.93$* |  |  |  |  |
| *Formula = Occurrence ~ Temperature\*Precipitation + Date + Region + Year + (1\|Unit)* |  |  |  |  |
| *Family = Binomial* |  |  |  |  |

<sup>a</sup>Significance codes: p < 0.001 = \*\*\*, p < 0.01 = \*\*, p < 0.05 = \*
<sup>b</sup>Variables were scaled prior to fitting

## Discussion

We confirm several trends reported or proposed elsewhere,^9,16,40,43^ and add clarity to relationships that have been more inconsistent,^18,35,36^ which are discussed further below. By analyzing data from colonies across Canada, we are also able to comment on prevalence of disease agents in some of the major beekeeping regions in this country; most notably, that *Varroa* and *M. plutonius* are more prevalent in BC than other provinces, at least within our sampling breadth, which was concentrated on two operations in the Lower Mainland area near Vancouver.

### Climatic effects

We found that temperature, wind, and precipitation were all influential factors for predicting *Varroa* occurrence, even after accounting for regional/operation and temporal effects, with temperature having a positive effect while wind and precipitation had a negative effect. That temperature has a positive impact above and beyond the effect of date aligns with expectations, as warmer temperatures promote brood nest expansion and foraging, giving the mites more opportunities to reproduce and possibly disperse. The negative effect of precipitation and wind are also intuitive, as such conditions limit bee flight (reducing dispersal opportunities), as well as possibly brood production via reduced foraging (reducing mite population growth rates).

Interestingly, Rowland *et al.*^9^ also found a positive effect of temperature and negative effects of wind and precipitation in an analysis of *Varroa* prevalence in the UK, which is consistent with our primary findings. However, our primary findings were only partially validated by our analysis of an independent *Varroa* dataset derived from similar regions in Canada.^47^ There, we found that temperature was the dominant climatic driver and the negative effect of wind was only observed among samples taken mid-season (June to early August), which is the sample point overlapping the most with the data in our initial analysis. Temperature and wind metrics had similar average magnitudes and spanned similar ranges in both the primary and validation datasets, so we expected to observe similar effects. We suspect that the different results are largely due to the two datasets being acquired over different time scales: while the 2020 and 2021 data were acquired from samples typically taken 2-4 weeks apart during the late spring and summer (with the exception of Ontario in 2020), the 2016 validation dataset spanned a considerably longer portion of the beekeeping season. It is sensible that the climatic variable promoting reproduction (an exponential process), rather than dispersal, would be most influential in this temporal scale.

Some research suggests that *Varroa* reproduction is inhibited when ambient temperatures surpass 35°C,^22,23^ and such conditions will likely become more common as the climate changes and the frequency of extreme heat events increases.^53–55^ Hot temperatures can also be harmful to honey bees,^7,56,57^ but not always.^58^ British Columbia and, to some extent, Alberta experienced a severe heat wave between June 25 and July 2, 2021,^59^ which falls within our study period. Temperatures reached >40°C in some parts of the Lower Mainland during this time, but the event was not strongly reflected in our data due to a combination of there being lower temperatures at our specific apiary locations and our process of computing three-week averages of daily means. Unfortunately, our sampling did not continue for long enough after the heat dome to measure any potential effect on *Varroa* reproduction.

Reported relationships between *Vairimorpha* spore intensities and temperature have been somewhat inconsistent, and our data add some clarity to the matter. A number of studies highlight broad seasonal patterns (with spore intensities decreasing between spring and fall)^18,19,21,32^, and some describe a negative relationship with temperature which is likely due to increased flight days,^18,35^ or possibly reduced ability to proliferate,^38^ though this is contradicted by Rangel *et al.*^36^ In our analysis, we found that temperature did not significantly predict a decline in spore intensities beyond the covarying effect of date, which supports the idea that flight days, and not temperature *per se,* is the limiting factor.

Further support of this reasoning is the interaction we observed between precipitation and wind **(Figure 4D**). These variables limit defecation flight opportunities independent of temperature, and while neither was sufficient to predict spore intensities on their own, precipitation did promote higher spore intensities under windier conditions. This makes sense given that the average total precipitation observed was not extreme (typically not more than 5 mm) and was therefore unlikely to sufficiently limit flights on its own. The likely crux of defecation flights as the operational factor could also be why Rangel *et al.* found no effect of temperature in their study of *Vairimorpha* in feral colonies in Texas. Those samples were collected in July over the course of 20 years, when one would expect the temperatures to be more than sufficient to enable flight days regardless of year-to-year deviations.

*M. plutonius* is often described as an opportunistic pathogen that is triggered to cause EFB disease based on a combination of nutritional deficiencies,^16,24,60^ pathogen genotype,^61,62^ and host genotype,^63^ among other factors. Given the possible connection with nutrition, it is conceivable that inopportune foraging conditions could promote disease and increase *M. plutonius* occurrence, but it has been suggested that strong nectar flows might actually also promote *M. plutonius* proliferation due to diversion of nursing care.^16,43^ While Wardell *et al.*^60^ states that no obvious climatic conditions account for disease incidence,^60^ this is not surprising for such a multifactorial issue, and others have postulated that climate may indeed play a role in part due to the strong regional differences that have been observed.^24,61^

We found both strong regional differences and a relationship with wind and precipitation, but no relationship with temperature after accounting for sampling date (**Figure 5**; **Table 6**). The highest occurrence frequencies were observed under both favourable (low wind, low precipitation) and unfavourable (high wind, high precipitation) conditions, with lower occurrence in intermediate zones. This pattern, while at first puzzling, may actually reflect both predictions: Increased foraging leads to a temporarily poorer brood care due to a shift from nursing to nectar-processing activities, while suboptimal foraging conditions also lead to poorer brood care via insufficient nutrition. It is also possible that favourable conditions lead to higher *M. plutonius* occurrence probability because this increases opportunities for dispersal, which is particularly relevant for this highly transmissible pathogen.^16^ Rowland *et al*. previously reported that high precipitation, but not wind, was positively linked to higher EFB incidence,^9^ and our results may differ in part because we analyzed *M. plutonius* occurrence, which is distinct from clinical diagnosis of EFB disease. The two metrics are different but nevertheless connected, especially as only 100-200 *M. plutonius* spores are required to cause disease.^42,43^ We expect that our method of PCR detection would be more sensitive to dispersal events in which the pathogen is transmitted but does not necessarily cause disease, which is consistent with our observation of higher occurrence under favourable climatic conditions.

### Regional effects

We observed strong regional effects for all three disease agents, with distinctly high incidence of *M. plutonius* and *Varroa* in BC, and variable levels of *Vairimorpha* spp. from year to year. However, we note that in our experiment we are unable to distinguish between effects of region and effects of beekeeping operation. We and others have previously observed an association between EFB and highbush blueberry pollination,^16,60,64^ and there is a high density of highbush blueberry fields in British Columbia’s Lower Mainland; however, two controlled field trials (including one derived from these data) do not support the notion that engaging in blueberry pollination causes *M. plutonius* infection.^45,65^ Therefore, perceived associations may actually be due to the seasonal timing of blueberry pollination. EFB outbreaks have also been observed in parts of Europe, where authorities responded with strict control measures including hive destruction and exclusion zones around affected apiaries.^66^ Especially given the high transmissibility of *M. plutonius* to neighboring colonies,^16,24,41^ strict control measures are essential for containment. BC has no provincial mandate for destroying or otherwise controlling affected colonies and apiaries, whereas in Ontario, where we observed zero *M. plutonius* detections, EFB is a reportable disease with enforceable movement restrictions and sanitation procedures.

British Columbia, at least in our sampling region of the Lower Mainland, also appears to suffer from high incidence of *Varroa* mites, particularly early in the season. This is despite colonies being treated for *Varroa* (in early April) before the beginning of the study. Although we did not formally test this, we speculate that this may be in part due to the region having notably mild winters, which lead to shorter brood breaks and earlier recommencement of brood rearing. These conditions mean that beekeepers have a shorter broodless window during which to apply a miticide treatment, and remaining mites are able to begin reproducing earlier. However, we note that the 2016 dataset contained yards in BC’s interior (the Thompson/Okanagan and Kootenay/Boundary regions), as well as the Lower Mainland and coastal regions, but the patterns observed there do not entirely agree with this idea. Based on average daily temperatures in the interior and Lower Mainland/coastal apiaries for 2016, we estimated that the spring brooding date in the interior was later, similar to the regions in other provinces. However, we still observed relatively high early-season *Varroa* incidence in the interior, with fitted probabilities of ∼0.5 (higher than all other regions). Therefore, factors yet unaccounted for are clearly at play, such as possibly beekeepers’ winter and early spring mite management procedures, and the timing of these in relation to when the experiment began.

### Conclusion

We present an analysis of common honey bee disease agents and how they vary according to regional, temporal, and climatic factors. Our results largely support patterns expected based on existing data and knowledge of how these disease agents reproduce and disperse. Nevertheless, confirmation of these trends is important on a nation-wide scale and across many climatic and ecotypes. Our analysis also provides additional insight into more ambiguous associations, particularly for how temperature affects *Vairimorpha* spore intensities and relationships between weather and *M. plutonius* occurrence. In most of our models, we observed a large degree of unexplained variation, suggesting that other factors, such as bee genetics, coinfections, or landscape factors may also influence pathogen and parasite dynamics, which were not investigated here. We hope that this and future studies will inform meaningful global predictive models of disease agents under future climate projections.

## Acknowledgements

We would like to acknowledge A. Achkanian, R. Bahreini, S. Bailey, V. Beger, C. Bryant, A. Chapman, M. Chihata, C. Currie, K. Dave, G.L. Ganesan, P. Grubiak, D. Holder, A. Ibrahim, M.A. Imrit, J. Janisse, J. Kearns, K. Lynch, D. Micholson, M. Munro, I. Ovinge, M. Peirson, K. Peters, Z. Rempel, R. Thygesen, D. Tran, A. Travas, and B. Vinson for their supportive efforts. We are also grateful to all members of the BeeCSI Consortium (https://beecsi.ca/beecsi-consortium) who participated this project.

This work was funded by Genome BC through the Genomic Innovation for Regenerative Agriculture, Food and Fisheries (GIRAFF) program. Data analyzed in this work was derived from the BeeCSI project, which was funded and supported by the Ontario Genomics Institute (OGI-185), Genome Canada (LSARP #16420), the Ontario Research Fund (LSARP #16420), Genome Quebec, and the Government of Canada through Agriculture and Agri-Food Canada (AAFC) Genomics Research and Development Initiative (GRDI) funding (AAFC J-002368). The work was also supported by the Canadian Honey Council and the Technology Transfer Program of the Ontario Beekeepers’ Association.

