## Supplementary Figures S1-S5 for "Climatic predictors of prominent honey bee (*Apis mellifera*) disease agents: *Varroa destructor*, *Melissococcus plutonius*, and *Vairimorpha* spp"

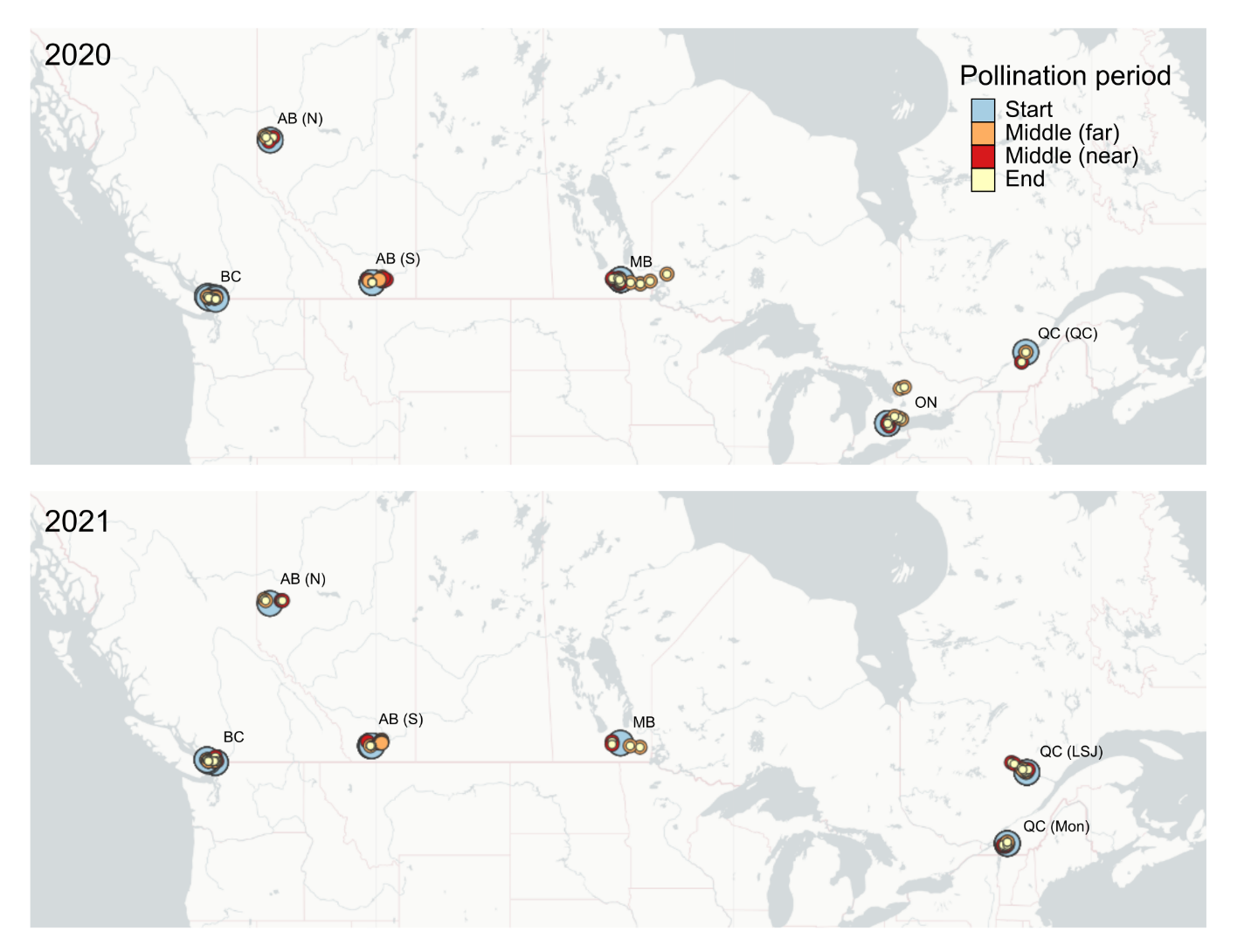

**Figure S1. Colony locations.** Each cluster of colony locations corresponds to one “region.” Colonies within regions originated from a single operation; therefore, regional and operational effects are indistinguishable. Colonies were sampled at three time points (at the start, middle, and end of the pollination period) with middle time points divided into groups of colonies near and far from focal crops (highbush blueberry and cranberry in BC, commodity canola in AB (N), commodity canola and canola seed in AB (S), commodity canola and soybean in MB, corn in ON, cranberry in QC (QC), apple in QC (Mon), and lowbush blueberry in QC (LSJ)).

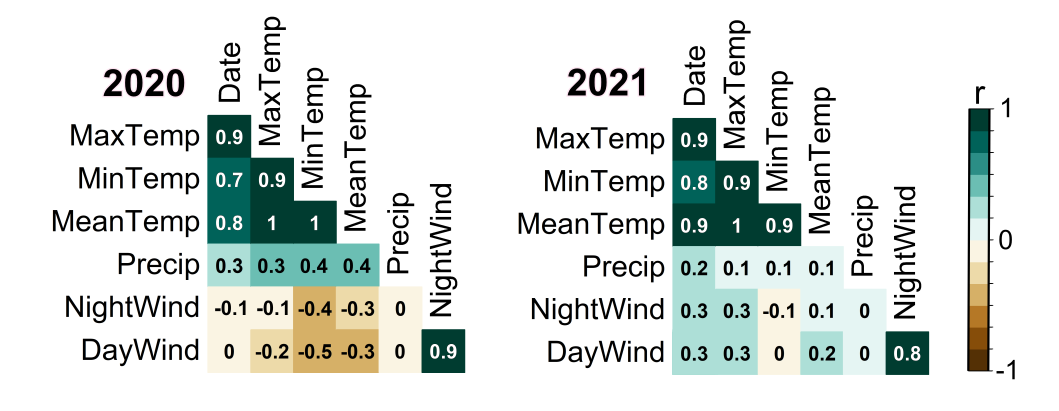

**Figure S2.** Pearson correlation matrix between climate variables. Numbers and colors indicate Pearson correlation coefficients.

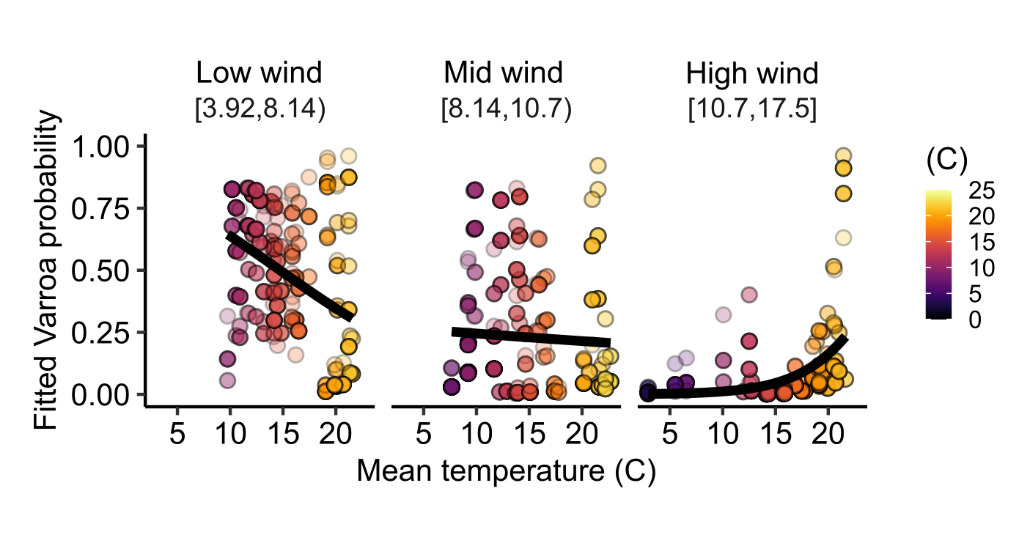

**Figure S3. Negative effect of wind speed on *Varroa* detections is moderated by high temperatures.** See **Table 2** for complete statistical reporting. Three equidensity wind speed bins are shown (km/h).

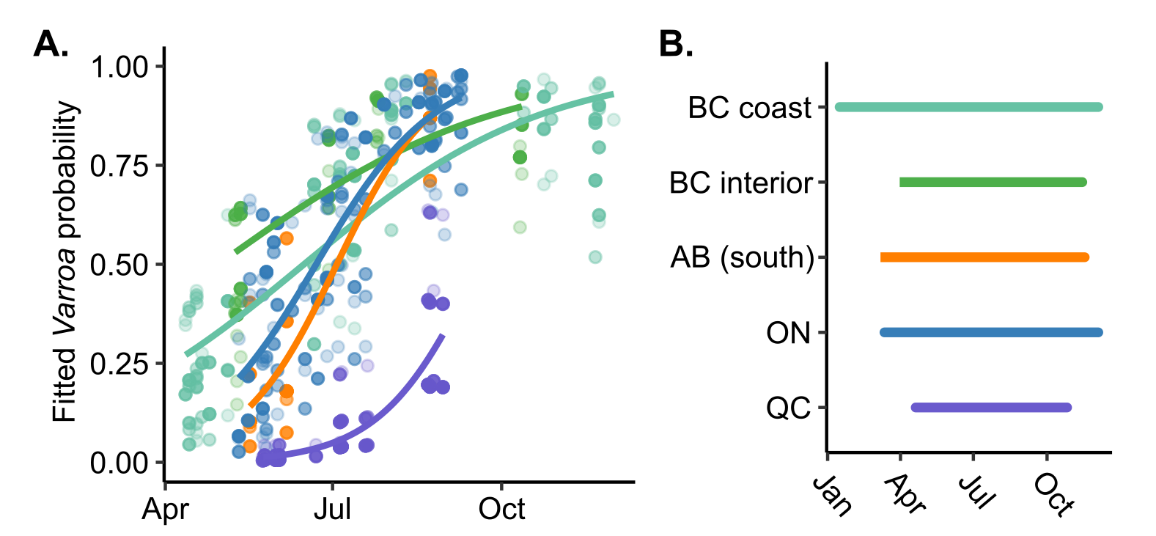

**Figure S4. Validation dataset: *Varroa* detection probability varies by location.** See **Table 3** for complete statistical reporting. The data modelled here are previously published^42^. A) Location differences in fitted *Varroa* detection probabilities. B) Length of brooding season in different locations. Defined as beginning on the date at which the seven-day sliding average temperature was at or above 5°C for at least three consecutive days, and ending on the date at which the seven-day sliding average temperature declined below 5 °C for ten or more days. Values were averaged across yards at each location.

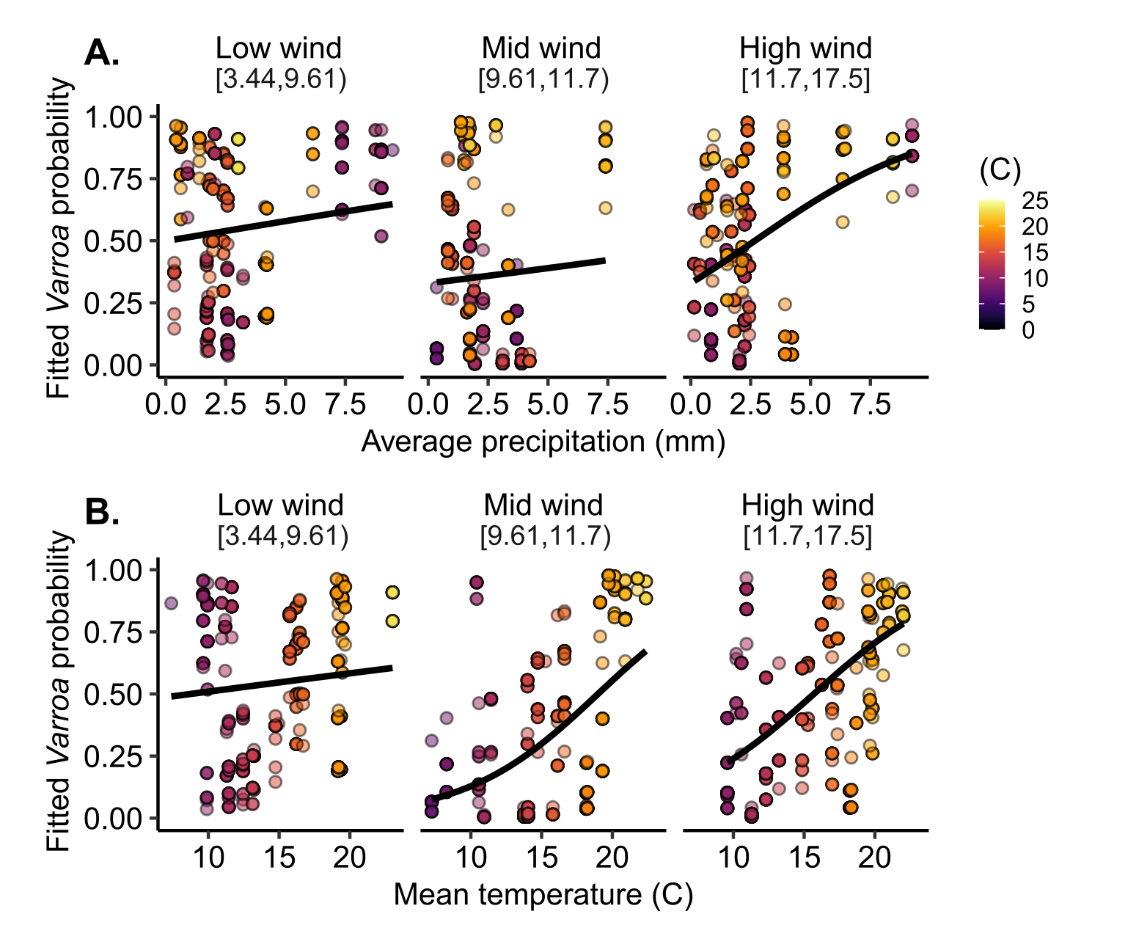

**Figure S5. Temperature, not wind, drives *Varroa* detection probability in long-term data.** *Varroa* detections derived from a 2016 validation dataset were fitted with a generalized linear mixed model with parameters and outcomes reported in **Table 3.** A) We found no significant effect of precipitation, which is inconsistent with the 2020 and 2021 data. B) Among other predictors, we found a significant interaction between temperature and wind. Temperature has a positive effect on *Varroa* detection frequency, as expected; however, this effect is exacerbated in mid-high wind conditions, which is inconsistent with the 2020 and 2021 data.
